# Turning a new leaf: PhenoVision provides leaf phenology data at the global scale

**DOI:** 10.1101/2025.09.26.678778

**Authors:** Erin L. Grady, Ellen G. Denny, Carrie E. Seltzer, John Deck, Daijiang Li, Russell Dinnage, Robert P. Guralnick

## Abstract

Plant phenology dictates many aspects of community function and ecosystem dynamics. Yet, global phenology data are still limited, especially in areas lacking monitoring programs. Here we present a new data resource, PhenoVision–Leaf, which extends a computer-vision pipeline utilizing iNaturalist digital image vouchers to produce global-scale leaf phenophase data for deciduous, woody genera. We first discuss our implementation of a new human annotation framework for leaf phenology on iNaturalist, aligning with phenophase definitions used by the larger phenology community. We then showcase the use of 165,988 crowdsourced annotated records to train a Vision Transformer model with a two-stage regime to maximize accuracy across single- and multi-image records. This approach extends Phenovision from scoring individual images to aggregating at the iNaturalist record level, better aligning with human annotation processes. Post-hoc validation showed high performance for detecting present green and colored leaves (>98% accuracy), and reasonable accuracy for breaking leaf buds (>87% accuracy). Applying PhenoVision–Leaf to over 26 million iNaturalist records yielded 5.6 million record-level phenology observations across 6,500 species and 57 families, filling geographic and taxonomic gaps. These data, now accessible through the Phenobase portal, establish a foundation for near real-time monitoring of leaf phenology, supporting global-scale synthesis analyses.

## INTRODUCTION

Phenology—the timing of life-cycle events like leafing, flowering, and fruiting—is a sensitive indicator of environmental change and a crucial driver of ecosystem dynamics (Caparros-Santiago et al., 2021). Despite a growing body of phenology research, significant data and theory gaps persist. While piecemeal advances have begun to close some gaps, phenology data remain limited in spatial, temporal, and taxonomic scope (Dinnage et al., 2025). Even in regions and countries where phenology is actively monitored, via amateur or professional networks, assembling these in-situ records into a coherent whole remains frustratingly hard (Stucky et al., 2018). Another potential source of phenology data at scale, remote sensing, can produce global estimates of the timing of leaf onset, peak greenness, and leaf senescence.

However, remote sensed phenology often cannot detect individual phenophases such as breaking leaf buds, flowering, and fruiting, and is a pixel-based measure rather than capturing phenology on individual whole plants (Diao et al., 2024). These many limitations ultimately restrict our ability to answer key questions in phenology and quantify broad-scale patterns of phenological change.

One promising path forward lies in leveraging community science platforms such as iNaturalist, which engages community members worldwide to generate millions of plant records monthly and continues to grow (Di Cecco et al., 2021). These records include images that often contain secondary data that can inform about phenology, such as evidence of the presence of certain phenological states. In recent work, we have shown the utility of computer-vision models for labeling the presence of flowers and fruits in field images (Dinnage et al., 2025), enabling scalable, high-quality phenology data production. Based on validation efforts, machine learning model accuracy, especially for flowers, often rivals or exceeds that of humans (Dinnage et al., 2025). Our work in Dinnage et al. (2025) was enabled both by the accessibility of iNaturalist occurrence metadata and images, and by the existence of a phenology annotation system already implemented in iNaturalist for flowers and fruits that provided extensive data useable for model training. Our model pipeline, PhenoVision, generated tens of millions of image-level machine labels about flowers and fruit presence, included rich metadata about how the modeling was performed, and was developed to seamlessly integrate with other existing phenology data (Dinnage et al., 2025). These data are included in a larger initiative, Phenobase (https://phenobase.org/), which encapsulates data integration processes and overall global phenology data services for the phenology community.

Here we showcase how we can extend the PhenoVision pipeline and Phenobase, both by producing global-scale leaf phenology information and by reconfiguring to produce record-level phenology data from iNaturalist field images and metadata. Unlike flower and fruit annotations, which were already available from iNaturalist for model training, leaf annotation training pipelines for PhenoVision had to start from scratch. We therefore describe the process in full for producing a high-quality, vast new leaf phenology dataset, capturing annotation for green leaves, colored leaves, and breaking leaf buds for woody, deciduous plants across the globe. We first begin by establishing and defining leaf phenophases that citizen scientists on iNaturalist would be able to capture, and then discuss implementing leaf phenology reporting on the platform with support from the iNaturalist team. We then describe decisions regarding implementation of our machine learning modeling frameworks, including improvements we have made in the pipeline, and moving from reporting annotations at the level of images to the iNaturalist record level.

The end result of the work presented here is a leaf phenology data resource that provides key information about the timing of green up and senescence for thousands of taxa, many of which have either never been available or have been available only for certain regions of the globe. We showcase how these new data can close spatial and taxonomic gaps, how we make this new data resource available to the community, and discuss next steps towards integrating phenology data resources in order to create Phenobase.

## METHODS

### Building leaf phenology annotations into iNaturalist

To build a large and reliable training dataset, we turned to the iNaturalist community to help classify leaf phenophases in community-collected photos of live plants. Prior to this effort, iNaturalist already included reproductive structure annotation capabilities (e.g., presence or absence of flowers and fruits), but did not yet have annotation options available for leaf phenophases. We felt confident that adding leaf annotation options to iNaturalist was a worthwhile next step, as iNaturalist users had proven to be successful at accurately classifying reproductive phenophases (Dinnage et al., 2025).

We selected four key leaf phenophases: breaking leaf buds, green leaves, colored leaves, and no live leaves. In collaboration with the iNaturalist team, we modified the USA National Phenology Network’s (USA-NPN) definitions to be clear and concise for a broad audience. In doing so, we largely retained the USA-NPN’s definitions while removing jargon (e.g., “petiole”) that might present as a barrier to making annotations by iNaturalist users (Table 1). This process resulted in definitions slightly less specific than those from the USA-NPN, which may leave more room for interpretation; however, we determined these choices worthwhile to engage a larger pool of iNaturalist users in the annotation process. Additionally, we made sure the selected terms and definitions aligned with terms included in the Plant Phenology Ontology (PPO) (Stucky et al., 2018), which is key for downstream integration with data from other sources (Table 1). The “no live leaves” category was included to help construct negative categories for each phenophase for training. Since absence cannot be determined from partial images of plants, we focused on detecting present phenophases only and did not explicitly train a model to detect “no live leaves”.

**Table 1.** Terms and their definitions for each of four annotation categories added to iNaturalist in comparison with the USA National Phenology Network (USA-NPN) and the Plant Phenology Ontology (PPO).

| iNaturalist |  | USA-NPN |  | PPO |  |
| --- | --- | --- | --- | --- | --- |
| Term | Definition | Term | Definition | Term | Definition |
| **Breaking leaf buds** | At least one bud with its first new leaf or needle not fully unfolded | **Breaking leaf buds** | One or more breaking leaf buds are visible on the plant. A leaf bud is considered "breaking" once a green leaf tip is visible at the end of the bud, but before the first leaf from the bud has unfolded to expose the leaf base at its point of attachment to the leaf stalk (petiole) or stem. | **Breaking vegetative bud** | A non-dormant vegetative bud that is in the bud burst stage. The bud burst stage is a shoot development stage during which a shoot system emerges from the bud. |
| **Green leaves** | At least one green leaf or needle (or normal healthy color such as reddish) | **Leaves\*** | One or more live, unfolded leaves are visible on the plant. A leaf is considered "unfolded" once its entire length has emerged from a breaking bud, stem node or growing stem tip, so that the leaf base is visible at its point of attachment to the leaf stalk (petiole) or stem. Do not include fully dried or dead leaves.<br>*Note that the USA-NPN "Leaves" phenophase includes "Colored leaves". The "Green leaves" iNaturalist definition most closely maps to USA-NPN's "Leaves" subtracting the "Colored leaves" subset. | **Non-senescent unfolded true leaf present** | An unfolded true leaf that is in the vascular leaf expanding unfolded stage (where the leaf has completed the process of unfolding but is still expanding in size) or the vascular leaf post-expansion stage (where the leaf is fully expanded but the vascular leaf senescent stage has not begun). |
| **Colored leaves** | At least one leaf or needle has late season or drought color | **Colored leaves** | One or more leaves show some of their typical late-season color, or yellow or brown due to drought or other stresses. Do not include small spots of color due to minor leaf damage, or dieback on branches that have broken. Do not include fully dried or dead leaves that remain on the plant. | **Senescing true leaf present** | An unfolded true leaf that is in the vascular leaf senescent stage. The vascular leaf senescent stage begins with the formation of an abscission zone at the base of a vascular leaf and ends with leaf separation and death. |
| **No live leaves** | No breaking leaf buds or green or colored leaves or needles are visible | NA |  | NA |  |

### Developing target taxa list

Because leaf phenophases are not as easy to detect and can vary in herbaceous and evergreen plants, we decided to train our leaf models only on deciduous, woody plant genera. We used a combination of the following three databases to capture key woody, deciduous genera: TRY (Kattge et al., 2020), BIEN (Maitner et al., 2018), and GIFT (Weigelt et al., 2020). This led to a list of 807 genera in total, including genera where all species are deciduous, but also those that have a mixture of deciduous and evergreen members. We used this list as a starting point and then filtered down further based on the following criteria: 1) We removed genera that are semideciduous since these are hard to diagnose regarding start of season; 2) We removed taxa where diagnosing breaking leaf buds is very challenging due to small size of leaves or difficulty applying the rubric for breaking buds; 3) We removed genera poorly represented on iNaturalist (fewer than 500 records) and a few others that exceeded this limit but had very limited numbers of visible breaking leaf buds or colored leaves. We did so because genera with very limited sampling are more challenging to correctly classify. These criteria resulted in a final list of 145 well sampled, diagnosable genera (Appendix S1; see Supporting Information with this article).

### Downloading training data and creating absent categories

iNaturalist records that included leaf phenophase annotations as of January 26, 2025 were downloaded for the list of target genera through a Darwin Core Archive prepared by the iNaturalist team for this purpose. This included annotations of green leaves, colored leaves, breaking leaf buds, and no live leaves. All images associated with these records were downloaded using iNaturalist’s open data repository on AWS (https://registry.opendata.aws/inaturalist-open-data/). For breaking leaf bud present annotations, we downloaded only images associated with records annotated by iNaturalist user @robgur, an expert on our team. We did so because initial assessment of breaking leaf bud annotations suggested that this was a harder task for annotators than other leaf annotation tasks. Given this, we decided it was worthwhile to reduce the error rate by utilizing a single expert who had performed a significant number of annotations across many taxa. The key tradeoff here is that we had a much smaller pool of breaking leaf bud present annotations to use in training.

We used these downloaded annotations and associated images to create absent categories, for model training purposes, for each of the three focal phenophases (green leaves, colored leaves, breaking leaf buds). We first assigned each image a value corresponding to each phenophase. The value for each phenophase was assigned as 1 if that phenophase was mentioned in the user annotations and 0 otherwise. If a phenophase was assigned a value of 0 for a certain image, meaning an iNaturalist user did not indicate that phenophase was present, that image was included in the absent category for that phenophase. It is important to note that this process was conducted only with images associated with at least one existing leaf phenophase annotation. The inclusion of the “no live leaves” phenophase in the downloaded annotations and associated images was critical, as it meant the absent categories included both images that had no leaves at all (e.g., no live leaves, green leaves absent) and images that had leaves in other phases, but not the focal phenophase (e.g., colored leaves present, green leaves absent). This process created three absence categories critical for model training: green leaves absent, colored leaves absent, and breaking leaf buds absent, and ensured representation in the data of all factorial combinations of the three categories.

### Quality of user-contributed leaf phenology annotations on iNaturalist

To understand the accuracy of iNaturalist user leaf phenophase annotations and the constructed absent categories, we conducted expert validation. On first look, we quickly noticed that breaking leaf bud annotations often included already unfolded leaves, falling outside our specific definition for this term. We therefore opted to train the breaking leaf bud model using internal expert annotations only, which is why we describe above that we only downloaded observation records associated with breaking leaf bud present annotations by @robgur. For the remaining categories, we validated 500 randomly selected images annotated for colored leaves present, 250 annotated for green leaves present, and 250 in each of our created absent categories (green leaves absent, colored leaves absent).

### Model training

The leaf phenology detection model follows a similar approach to that described in detail in Dinnage et al. (2025), with several key modifications. We used a Vision Transformer (ViT) Large architecture with 16×16 patches (Dosovitskiy et al., 2020), leveraging the same masked autoencoder (MAE) pretraining approach (He et al., 2022). Rather than starting from the PlantCLEF “virtual taxonomist” pretrained model as in the original PhenoVision work, we initialized our model with the trained PhenoVision flower and fruit detection model (Dinnage et al., 2025), providing an additional layer of phenology-specific pretraining. The model architecture was adapted to output three binary classifications corresponding to our leaf phenophases: green leaves, colored leaves, and breaking leaf buds. We replaced the final classification layer with a new randomly initialized linear layer that transforms the 1024-dimensional latent representation into a three-dimensional vector, passed through a sigmoid activation function to produce probabilities between 0 and 1 for each phenophase. Before training, we split our annotated dataset into 80% for training, 10% for validation, and 10% for testing to ensure robust model evaluation and prevent overfitting.

A critical challenge in training was that iNaturalist phenology annotations are associated with whole observations rather than individual images, and observations can contain multiple images, sometimes showing different parts of the plant. This creates ambiguity about which specific images actually display the annotated phenophases. To address this, we employed a two-stage training regime designed to maximize the use of available data while ensuring annotation accuracy. In the first stage, we trained exclusively on observations containing single images, guaranteeing direct correspondence between annotations and image content. To address extreme class imbalance, we separated training data into two groups: observations with only green leaves present (the most common or “background” state), and observations with any other phenophase combination. We then upsampled the non-background observations with replacement to match the number of background observations, creating a more balanced dataset that was fixed for all training epochs. The same balancing procedure was applied to the validation set. Note that this creates duplicate data but any impact of such duplication is minimized because we used image augmentation which made sure each duplicate had different random augmentations. Training used a batch size of 384 with layer-wise learning rate decay (layer_decay = 0.65), AdamW optimization, and enhanced data augmentation including RandAugment (auto_augment = ‘rand-m9-mstd0.5-inc1’), as implemented in the ‘timm’ Python package. This applies a random image transformation from the set of AutoContrast, Equalize, Rotate, Posterize, Solarize, Color adjustment, Contrast, Brightness, Sharpness, and geometric transformations (Shear, Translate), as well as Random erasing with pixel-level replacement. The model was trained for 4 epochs.

The second stage employed a semi-supervised approach to incorporate multi-image observations. We used the first-stage model to predict phenophases for all images from multi-image observations, then retained only those images where: (1) the model predictions exceeded 0.95 confidence, and (2) the predicted phenophases matched the user’s original annotations. This approach automatically identified which images from multi-image observations actually displayed the annotated structures. These validated images were combined with the original single-image training data, and the model was fine-tuned for an additional 4 epochs. This two-stage process maximized training data utilization while maintaining high confidence in the correspondence between images and their phenological labels. The model was implemented using PyTorch with the timm and transformers libraries, integrated with R preprocessing via reticulate, and trained on a single NVIDIA A100 GPU.

### Determining model threshold values and equivocal buffer zones

The trained model outputs continuous probability values between 0 and 1 for each leaf phenophase, requiring threshold values to convert these probabilities into binary presence/absence predictions. To optimize these decision boundaries, we followed the approach detailed in Dinnage et al. (2025). We optimized detection threshold values and equivocal buffer zones for each of the three leaf phenophases independently using the held-out validation dataset. We evaluated performance across a range of threshold values, optimizing for maximum True Skill Statistic (or TSS which balances sensitivity and specificity). After establishing optimal detection thresholds for green leaves, colored leaves, and breaking leaf buds, we empirically determined equivocal buffer zones by examining model accuracy across binned prediction values (100 bins from 0 to 1), determining when a moving average first equaled or exceeded 75% accuracy. Images with model predictions falling within these equivocal zones were classified as equivocal detections and excluded from the final dataset, while those outside these zones were classified as unequivocal. This approach maximized annotation quality by retaining only unequivocal predictions while preserving the majority of high-confidence detections. These optimized thresholds and buffer zones were saved and applied consistently across all subsequent model predictions.

### Model validation and machine-labeling

We first validated using the held-out test set and then downloaded and machine-labeled a subset of randomly selected iNaturalist images for additional expert validation. We expert validated 500 images machine-labeled as detected, 500 labeled as undetected, and 250 labeled as equivocal in each of the three categories (green leaves, colored leaves, breaking leaf buds). We focused on model accuracy for unequivocal detection, since for final downstream data, we only retained images unequivocally labeled as detected. This is because images are often not of the entire plant and we therefore cannot confirm phenophase absence. We then downloaded all iNaturalist images linked to records of our target taxa uploaded before May 27, 2025. Each image was machine-annotated for green leaves, colored leaves, and breaking leaf buds. Each machine annotation was designated as equivocal or unequivocal based on whether the model’s output value for a particular trait fell within its equivocal buffer zone as determined by the procedure described in the previous section.

### Producing record-level phenology information

After conducting machine annotation at the image level, the next step was to aggregate these images to make a decision about the presence of a phenophase at the iNaturalist record level. This step was critical for correct use of these data downstream. For each record, we counted the number of images in each of the machine-labeled categories (detected-unequivocal, undetected-unequivocal, equivocal) for each phenophase. We then filtered out all categories except detected-unequivocal. As long as at least one image in a record had a phenophase detected unequivocally, that record will appear in the data as “present” for that phenophase. This resulted in a final dataset that included only unequivocal “present” observations at the record x phenophase level.

### Evaluating increases in spatial and taxonomic coverage

To evaluate increases in data coverage resulting from the machine labeling of these data in comparison to existing iNaturalist user leaf phenophase annotations, we followed methods described in Dinnage et al. (2025). To visualize increases in taxonomic data coverage, we determined the proportion of leaf phenology records (for each phenophase) contributed by human-annotators versus machine-labeled in each of the 145 key genera. We then used 100 km x 100 km equal-area grid cells plotted globally using the R package sf (Pebesma, 2018) to identify cells with previously no phenology information that now have leaf data produced through the PhenoVision–Leaf pipeline, plot the total number of records available across space, and visualize the proportion of records in each cell contributed by PhenoVision–Leaf.

## RESULTS

### Downloading training data and creating absent categories

We downloaded 165,988 total iNaturalist records for training, which included 326,128 images. 53% (88,184) of these records only had one image and were used in the first round of training and 47% (77,804) had multiple images and were used in the second round of training (Appendix S2).

### Quality of user-contributed leaf phenology annotations on iNaturalist

Expert validation found that user-contributed annotations were highly accurate for green leaves, and moderately accurate for colored leaves present and for the colored leaves constructed absent class (Table 2). For the colored leaves present category, we found that iNaturalist users sometimes mistakenly annotated fully senesced leaves and leaves where the healthy color is red as “colored leaves present”. Breaking leaf bud annotations from all users were not formally validated, as we deemed them en masse inaccurate on first pass and decided to use only expert-annotated records for breaking leaf bud training. The constructed absent class for green leaves was less accurate (Table 2). This is likely due to records annotated on iNaturalist as “colored leaves present” or “breaking leaf buds present” that also include green leaves but lack a “green leaf present” annotation. Additional problems arise from confusion about what constitutes a breaking leaf bud, as some images annotated only as “breaking leaf buds present” actually show buds that have already opened and should be annotated as “green leaves present”. As a result, some records were included in the training dataset as “green leaves absent” even though green leaves are present in the image(s). However, since the PhenoVision pipeline focuses on present-only observations downstream, model tuning is focused on maximizing present accuracy and absent class accuracy is less critical for training.

**Table 2.** Expert validation results for user-contributed leaf annotations on iNaturalist. Absent classes were built using images that had at least one leaf phenophase annotation, but had no annotation for the focal phenophase. Bolded numbers represent cases of agreement between iNaturalist community annotators and expert annotators.

| Expert annotation | Green leaves present (250) | Green leaves absent (250) | Colored leaves present (497) | Colored leaves absent (250) |
| --- | --- | --- | --- | --- |
| Present | **99.6%** (249) | 24.4% (61) | **92%** (457) | 8.8% (22) |
| Absent | 0.4% (1) | **72.4%** (181) | 7.2% (36) | **91.2%** (228) |
| Unknown | 0.0% (0) | 3.2% (8) | 0.8% (4) | 0.0% (0) |

### Model validation

Expert validation found the PhenoVision–Leaf models to be highly accurate for detecting present green leaves (98.6% accuracy) and present colored leaves (99.4% accuracy), and moderately accurate for detecting present breaking leaf buds (87%) (Table 3). Model accuracy for detecting breaking leaf buds likely reflects the difficulty of the task, given it is a transitory state and more uncommon than either green leaves or colored leaves. It also reflects that there were far less training data available for model calibration since all leaf buds annotations were generated by one person. We also note the very high success of labeling unequivocal colored leaf presence given training input quality, which we attribute to the filtering process which removes equivocal machine annotations. Compared to expert validation results, validation on the held-out test data demonstrated higher accuracy for detecting present green leaves (99.3%), lower (although still relatively high) accuracy for detecting present colored leaves (91.2%), and considerably higher accuracy for detecting breaking leaf buds (98.9) (Appendix S3). However, as noted above, iNaturalist users sometimes struggle to correctly classify colored leaves (8% error rate) and breaking leaf buds. Since held-out data validation is based on the annotations of iNaturalist users, these accuracy values may be inflated or deflated.

**Table 3.** Validation results for PhenoVision-produced green leaves, colored leaves, and breaking leaf buds labels. 500 images were validated in the detected (Det.) and undetected (Undet.) categories and 250 in the equivocal (Equiv.) category by an expert human annotator. Bolded numbers represent cases of agreement between the model and expert annotators. Shaded cells represent accuracy scores for data retained and included in downstream Phenobase data.

| Expert | Green leaves |  |  | Colored leaves |  |  | Breaking leaf buds |  |  |
| --- | --- | --- | --- | --- | --- | --- | --- | --- | --- |
|  | Det. | Undet. | Equiv. | Det. | Undet. | Equiv. | Det. | Undet. | Equiv |
| Present | **98.6%**<br>(493) | 12.8%<br>(64) | 59.6%<br>(149) | **99.4%**<br>(497) | 7.2%<br>(36) | 62.8%<br>(157) | **87%**<br>(435) | 1.6%<br>(8) | 7.6%<br>(19) |
| Absent | 1.2%<br>(6) | **82.6%**<br>(413) | 30.8%<br>(77) | 0.6%<br>(3) | **91.8%**<br>(459) | 28.8%<br>(72) | 9.4%<br>(47) | **96.6%**<br>(483) | 83.2%<br>(208) |
| Unknown | 0.2%<br>(10) | 4.6%<br>(23) | 9.6%<br>(24) | 0.0%<br>(0) | 1.0%<br>(5) | 8.4%<br>(21) | 3.6%<br>(18) | 1.8%<br>(9) | 9.2%<br>(23) |

### Model errors

During human expert validation, we identified false positive errors for each phenophase, meaning the model detected a phenophase that doesn’t actually occur in the image (Figure 1). For green leaves the false positive rate was 1.2%, for colored leaves 0.6%, and for breaking leaf buds 9.4%. Here, we focus on false positive errors, as these are the only errors that occur in our final dataset given that we only reported detected results in the data product. Green leaves and colored leaves had very low false positive error rates, with background leaves causing the majority of errors for green leaves and leaf color not due to seasonal senescence or drought causing the majority of errors for colored leaves (Figure 1). As expected, breaking leaf buds had a higher false positive rate, with structures of certain taxa and abnormalities such as broken stems causing model confusion (Figure 1).

**Figure 1.**
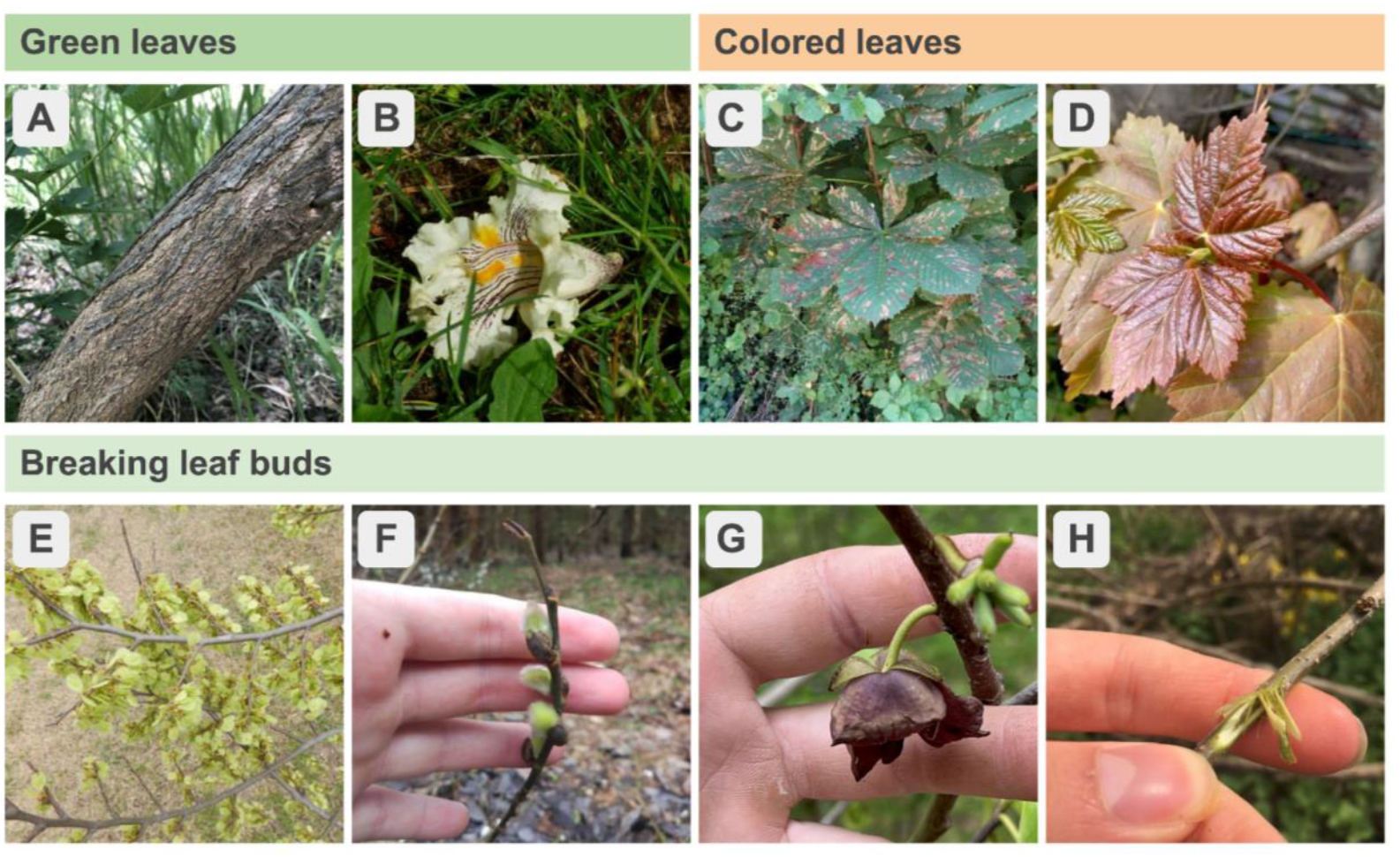
Examples of false positives, images where the model reports a phenophase as “present” when it is actually “absent”, for green leaves, colored leaves, and breaking leaf buds. *Green leaves*: (A) The only visible part of the target organism in this image is the stem. The model likely reported green leaves as “present” because green leaves occur in the background. (B) The only visible part of the target organism is a detached flower. The model likely reported green leaves as “present” because of the surrounding green leaves. *Colored leaves*: (C) While the leaves are colored, this is not due to seasonal change. (D) The reddish leaves in this image represent new, healthy growth and not seasonal senescence. *Breaking leaf buds:* (E) The presence of elm (*Ulmus* sp.) fruits in this image, with their unique shape, likely confuse the model. (F) Willow (*Salix* sp.) The presence of flower buds, which have a shape similar to leaf buds, likely confuse the model. (G) Pawpaw (*Asimina* sp.) often confuse the breaking leaf bud model, likely due to the sepals. (H) In this image, a broken stem tricks the model. Images courtesy of iNaturalist users: A-richkiker, B-mhalsted, C-bobwardell, D-sezzylou, E-bob777, F-sansan_94, G-axicus, H-mws.

### Downloading and machine-labeling images

We downloaded and labeled a total of 26,448,560 iNaturalist records for the target taxa, which included 43,610,500 images, observed by May 27, 2025 (Table 4). 64.3% of these records had just one associated image, and the mean and median number of images per record were 1.65 and 1, respectively.

**Table 4.** Final results for the number of images and records for each phenophase machine-labeled as detected, undetected, and equivocal. Shaded cells represent data retained and included in downstream Phenobase data.

| Machine annotation | Green leaves |  | Colored leaves |  | Breaking leaf buds |  |
| --- | --- | --- | --- | --- | --- | --- |
|  | Images | Records | Images | Records | Images | Records |
| Detected | 8,690,519 | 5,201,120 | 344,645 | 244,747 | 215,463 | 151,240 |
| Undetected | 1,643,014 | 963,913 | 9,653,435 | 5,612,048 | 10,409,882 | 6,022,125 |
| Equivocal | 569,092 | 469,023 | 904,545 | 701,477 | 277,280 | 225,300 |

### Increases in spatial and taxonomic coverage

In total, the final PhenoVision–Leaf phenology dataset includes 5,597,107 new phenology records, 5,201,120 of green leaves, 244,747 of colored leaves, and 151,240 of breaking leaf buds generated from machine annotation of iNaturalist records. These records have standardized fields, including key components such as day of year, location, taxon, phenophase, quality metrics, and a model identifier. They are mapped to the Plant Phenology Ontology (PPO), making them easily integratable with other data sources. The final mapped dataset is included in the open-access Phenobase web portal (https://phenobase.netlify.app/), along with flower and fruit iNaturalist records, observations from the USA National Phenology Network and other monitoring networks, and flower records from herbarium specimens from GBIF. See data and code availability below for more details.

These data represent 6,501 species from 145 genera and 57 plant families (Figure 2). For records of green leaves, the mean and median number of machine-annotated records per genus were 35,869 and 16,443. For records of colored leaves, the mean and median number of machine-annotated records per genus were 1,700 and 509. For breaking leaf bud records, the mean and median number of machine-annotated records per genus were 1,043 and 350.

**Figure 2.**
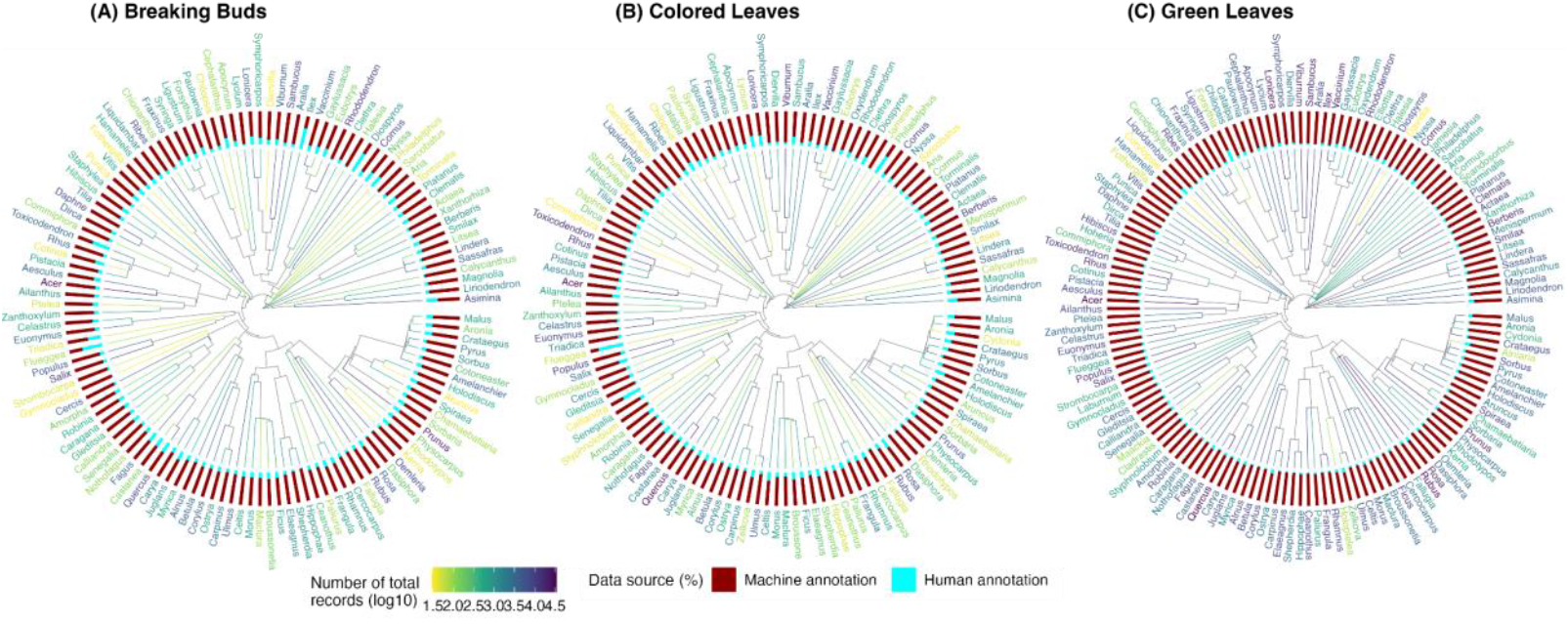
Machine-labeled leaf data vastly increases the amount and breadth of phenology data available across deciduous, woody taxa. Each of the 145 tips represents a plant genus on the target taxa list. Colors on the phylogeny branches represent the total number of leaf phenology observations available for that taxon (from iNat annotations and PhenoVision annotations). Bars demonstrate the proportion of machine-annotated records (dark red) versus the proportion of human-annotated records (cyan) available for each genus. For most genera, the vast majority of available phenology records are those machine-labeled through our PhenoVision pipeline.

Globally, there are 8,515 100km x 100km grid cells that have at least one leaf phenology record contributed by PhenoVision–Leaf (Figure 3). Breaking leaf bud information was available for 3,394 cells, with 1,667 of these cells that did not have any iNaturalist user-annotated records previously. Colored leaf information was available for 3,745 cells, with 1,811 cells that did not have any human-annotated records. Green leaf records were available for 8,489 cells, and 4,342 of them did not have human-annotated records. Across grid cells, the mean proportions for machine-annotated records were 74.9% for breaking buds, 74.7% for colored leaves, and 88.9% for green leaves. After removing cells with fewer than 10 records, there are 1,348 cells left for breaking buds, 1,719 cells for colored leaves, and 5,045 cells for green leaves (Appendix S4).

**Figure 3.**
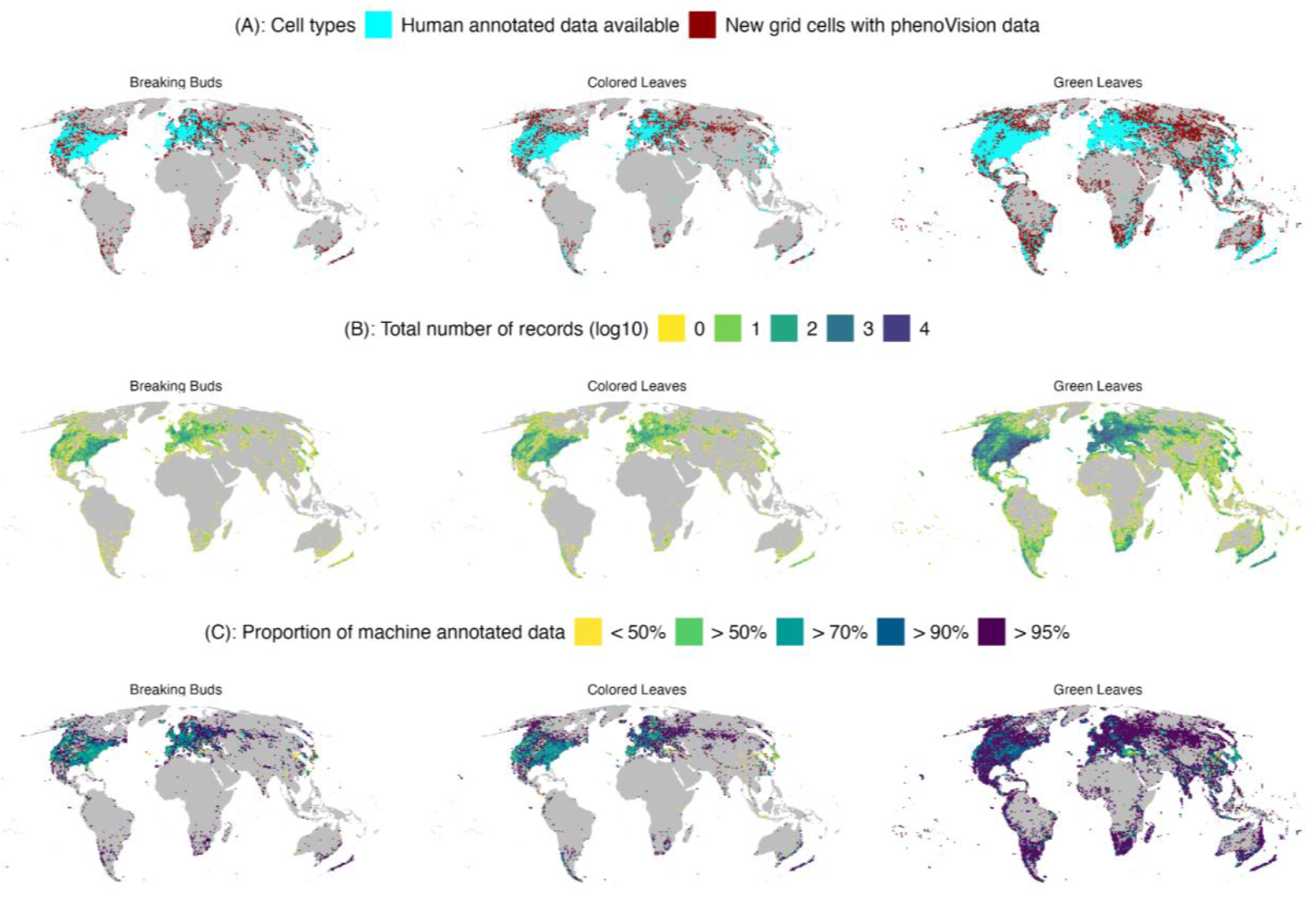
Global spatial coverage of breaking leaf bud, colored leaf, and green leaf phenology records at a 100km x 100km equal-area grid cell level. Each colored grid cell represents a cell with at least one phenophase record; grey cells represent gaps. (A) The spatial distribution of grid cells with at least one manually-annotated phenophase present record (light blue) and new grid cells for which PhenoVision provided machine-annotated data (dark red). (B) The spatial distribution of record counts (log_10_-transformed) for each leaf phenophase generated by PhenoVision. (C) The spatial distribution of the proportion of phenology data generated by PhenoVision.

## DISCUSSION

Here, we present a new and extensive global leaf phenophase dataset and a significant update to our PhenoVision pipeline, a workflow developed to automate phenophase annotations of community-collected iNaturalist records. Previous work of ours (Dinnage et al., 2025) automated image-level annotation for flowers and fruits. In this work, we add machine labeling of leaf phenophases and the ability to aggregate to the iNaturalist record level, unlocking millions of new phenology records for green leaves, colored leaves, and breaking leaf buds and helping lay the groundwork for near real-time, global phenology monitoring using iNaturalist. These annotations are generally highly accurate, fill spatial and taxonomic data gaps, and have immediate applications for answering fundamental large-scale phenology questions. Phenology records produced through the PhenoVision pipeline, for both reproductive and leaf phenophases, are designed for integration with other sources of phenology information, such as in situ monitoring programs and herbarium specimens, and for inclusion in Phenobase, an open, accessible and integrated data resource for the phenology research community.

Our collaboration with iNaturalist allowed us to rely on crowdsourced annotations to train the leaf phenophase models for green leaves and colored leaves. After introducing the leaf annotation tools on the iNaturalist platform, we led a phenology-focused webinar for the iNaturalist community, serving as an introduction to the concept of phenology, providing examples of phenology research enabled by iNaturalist, and demonstrating how to add annotations to pre-existing observations. To bring home just how quickly we moved from providing a means for iNaturalist participants to label leaf phenophases to producing high quality models, iNaturalist launched leaf annotation tools in June 2024 and we had models for colored leaves and green leaves developed by Spring 2025. This success can be attributed to the hard work of thousands of iNaturalist community volunteers who worked to provide annotations across many taxa.

While the crowdsourcing approach worked nicely to generate training data for both green and colored leaf categories, our initial review of breaking leaf bud labels suggested that this task is much harder. We found lower accuracy for human labels of breaking leaf buds, suggesting that it is more difficult to classify more ambiguous or transitional stages (e.g. breaking leaf buds) and that these tasks may be better suited to trained volunteers through initiatives like the USA National Phenology Network (Crimmins et al., 2022). A takeaway is that no single strategy is universally optimal and the approach should be tailored to the complexity of the task, expertise of the annotators, and downstream data needs. Here, our decision to use only expert annotations for breaking leaf buds allowed for a more controlled annotation process and higher annotation accuracy, but there are still trade-offs, especially with the quantity of annotations for training that can be generated this way. These trade-offs may partially explain why we see lower accuracy for the breaking leaf bud model than for other phenophases, indicating that even with careful consideration and high-quality training data, some annotation tasks are either simply more challenging than others or require significantly more human labeling data to achieve higher accuracy.

Despite not yet achieving the highest precision possible for breaking leaf buds, in general our approach demonstrates that machine vision models can produce high-quality results even when trained on relatively limited pools of human-annotated data. In our original Phenovision work (Dinnage et al., 2025), we had far more training data for flowers and fruits – by a factor of over 10 – compared to the leaf data here. Such large training sets are not always practical or possible to generate. Data here are of a size more typical for individual research groups, and it is encouraging that models can still produce research-ready data even with smaller training sets. The machine learning methods used here, including pretraining, class balancing, and data augmentation likely made this possible and are more crucial in these cases of limited data.

Beyond the new corpus of leaf phenology annotations, a key component of this work was updating our pipeline to report record-level, rather than simply image-level, annotations. This represents a key step in extracting and delivering information collected across the globe by iNaturalist observers to the phenology research community in a timely and scalable way. iNaturalist records reflect a process in which an observer records and uploads an instance of an organism, and photo evidence of this instance can include multiple images wrapped within one record. Thus, human labeling of phenophases on iNaturalist is at the record level, not per field image. While classifying phenophases at the image-level has utility of its own, aggregating our machine annotations to the iNaturalist record level provides research-ready and user-friendly data aligned with the human observation and annotation processes. We plan to update our pipeline to present record-level phenology information not just for leaf phenophases, but for reproductive phenophases as well (Dinnage et al., 2025). With iNaturalist users often making tens of thousands of flowering plant observations per day, our pipeline to quickly annotate these records for both leaf and reproductive plant phenophases enables a new kind of phenology monitoring on a global scale.

As evidenced in the above phylogenetic and spatial coverage plots (Figures 2 and 3), these models unlock data across diverse taxa and regions where phenology knowledge has previously been sparse or absent. We point especially to areas, such as remote areas of Eurasia including temperate forests and more boreal and arctic habitats, that, to our knowledge, have limited openly available phenology data. We also emphasize the broad swath of taxa for which phenology data are now available, many of which have never been part of global monitoring programs, which often focus on a narrow subset of canopy or subcanopy trees for woody deciduous taxa. Our goal here was not only to address these gaps, but also to make the resulting data readily accessible for the greater phenology research community. With increased access to phenology data at broad spatial and taxonomic scales, researchers can make strides to address fundamental questions about the impacts of global change on vegetation dynamics, including biomass production, carbon and nitrogen cycling, and general ecosystem functioning. In particular, data from more remote areas and especially those facing the most rapid changes, such as higher latitude areas in Eastern Europe and into Asia, and the ability to monitor across the broadest environmental gradients both hold promise for understanding the magnitude of phenological response (Brown et al., 2012). In turn, these can help us better understand species resilience in the face of continuing change.

These data are part of a larger effort, Phenobase, to integrate and provide openly available, global phenology data from multiple sources, ranging from purely incidental data such as iNaturalist and historical specimen data, to more structured monitoring data provided by programs such as the USA National Phenology Network and other regional monitoring programs. With this goal in mind, we have been deliberate in standardizing data fields and in clearly communicating the strengths and limitations of our models, through fields depicting phenophase and family-specific model accuracy, to facilitate interoperability and trust. We are also working on a set of companion guides so that end users are best able to use Phenobase resources most successfully. Ultimately, our aim is to build not just data resources but the community capacity to work towards understanding global-scale changes in phenology and their impacts on species and ecosystems.

## Acknowledgements

The authors would like to particularly acknowledge the members of the iNaturalist community who provided phenology annotations for training, and those who collected records and photo vouchers used throughout this process. They also extend thanks to the iNaturalist team for their help implementing and promoting leaf annotation capability on the platform. This work was funded by the National Science Foundation, in particular grants DBI2223512 to Robert Guralnick and DBI2223508 to Daijiang Li.

## Author Contributions

Robert Guralnick and Russell Dinnage conceived the ideas and designed methodology with input from Erin Grady, Daijiang Li, Carrie Seltzer, Ellen Denny, Ramona Walls and John Deck. Russell Dinnage developed the machine learning model and workflow with input from Robert Guralnick, Erin Grady and Daijiang Li. Daijiang Li downloaded data from iNaturalist with help from Carrie Seltzer. Erin Grady and Robert Guralnick led the model validation effort; Erin Grady, Russell Dinnage, Robert Guralnick and Daijiang Li analysed the data. Erin Grady, Robert Guralnick, and Russell Dinnage led the writing of the manuscript. All authors contributed critically to the drafts and gave final approval for publication.

## Data Availability Statement

Full code related to this manuscript is publicly available on GitHub (https://github.com/Phenobase/phenovision). The fully trained Phenovision–Leaf model is available on Hugging Face (https://huggingface.co/phenobase/phenovisionL). A dataset with record-level machine annotated leaf phenology data (present annotations only) is available on Zenodo (https://doi.org/10.5281/zenodo.17107251). These records are also available on the PhenoBase data portal (https://phenobase.netlify.app/), where they are integrated with in situ phenology observations and phenology information from herbarium specimens.

## Supporting Information

Additional Supporting Information may be found online in the Supporting Information section at the end of the article.

Appendix S1. List of included woody, deciduous genera.

Appendix S2. Number of images included in each category for each round of training. Appendix S3. Validation on held-out test data.

Appendix S4. Global spatial coverage of breaking leaf bud, colored leaf, and green leaf phenology records at a 100km x 100km equal-area grid cell level.

