## Appendix S2 for "Turning a new leaf: PhenoVision provides leaf phenology data at the global scale"

**Table S2.** Number of images included in each category for each round of training.

| **Training round** | **Green leaves present** | **Green leaves absent** | **Colored leaves present** | **Colored leaves absent** | **Breaking leaf buds present** | **Breaking leaf buds absent** |
| --- | --- | --- | --- | --- | --- | --- |
| Round 1 | 77, 270 | 10,914 | 8,471 | 79,713 | 4,479 | 83,705 |
| Round 2 | 205,261 | 32,683 | 30,373 | 207,571 | 44 | 237,900 |
