## Appendix S3 for "Turning a new leaf: PhenoVision provides leaf phenology data at the global scale"

**Table S3.** Validation on held-out test data. User annotations represent those made by iNaturalist users. Percentages for detected and undetected represent the percentage of images annotated as present or absent that the model did or did not detect. “Equivocal” values represent the percentage of images classified as equivocal. Shaded calls represent model-annotator agreement for present annotations included in the final dataset. The present, equivocal category is bolded and represents how many present images were “lost” from the dataset. We included No Live Leaves for evaluation of model performance although these were not included in the final datasets made available.

| **User Annotation** | **Green leaves** | | | **Colored Leaves** | | | **Breaking Leaf Buds** | | |
| --- | --- | --- | --- | --- | --- | --- | --- | --- | --- |
|  | **Det.** | **Undet.** | **Equiv.** | **Det.** | **Undet.** | **Equiv.** | **Det.** | **Undet.** | **Equiv** |
| **Present** | 99.3% (9323) | 0.7% (68) | **2.4% (228)** | 91.2% (640) | 8.8% (62) | **34.6% (371)** | 98.9% (536) | 1.1% (6) | **7.2% (42)** |
| **Absent** | 3.3% (42) | 96.7% (1242) | 8.3% (117) | 0.1% (10) | 99.9% (9396) | 0.1% (10) | 0.2% (23) | 99.8% (10368) | 0.4% (45) |

| **User Annotation** | **No Live Leaves** | | |
| --- | --- | --- | --- |
|  | **Det.** | **Undet.** | **Equiv** |
| **Present** | 96.5% (329) | 3.5% (12) | **9.1% (34)** |
| **Absent** | 0.2% (348) | 99.8% (9473) | 10.8% (1187) |
