## Appendix S4 for "Turning a new leaf: PhenoVision provides leaf phenology data at the global scale"

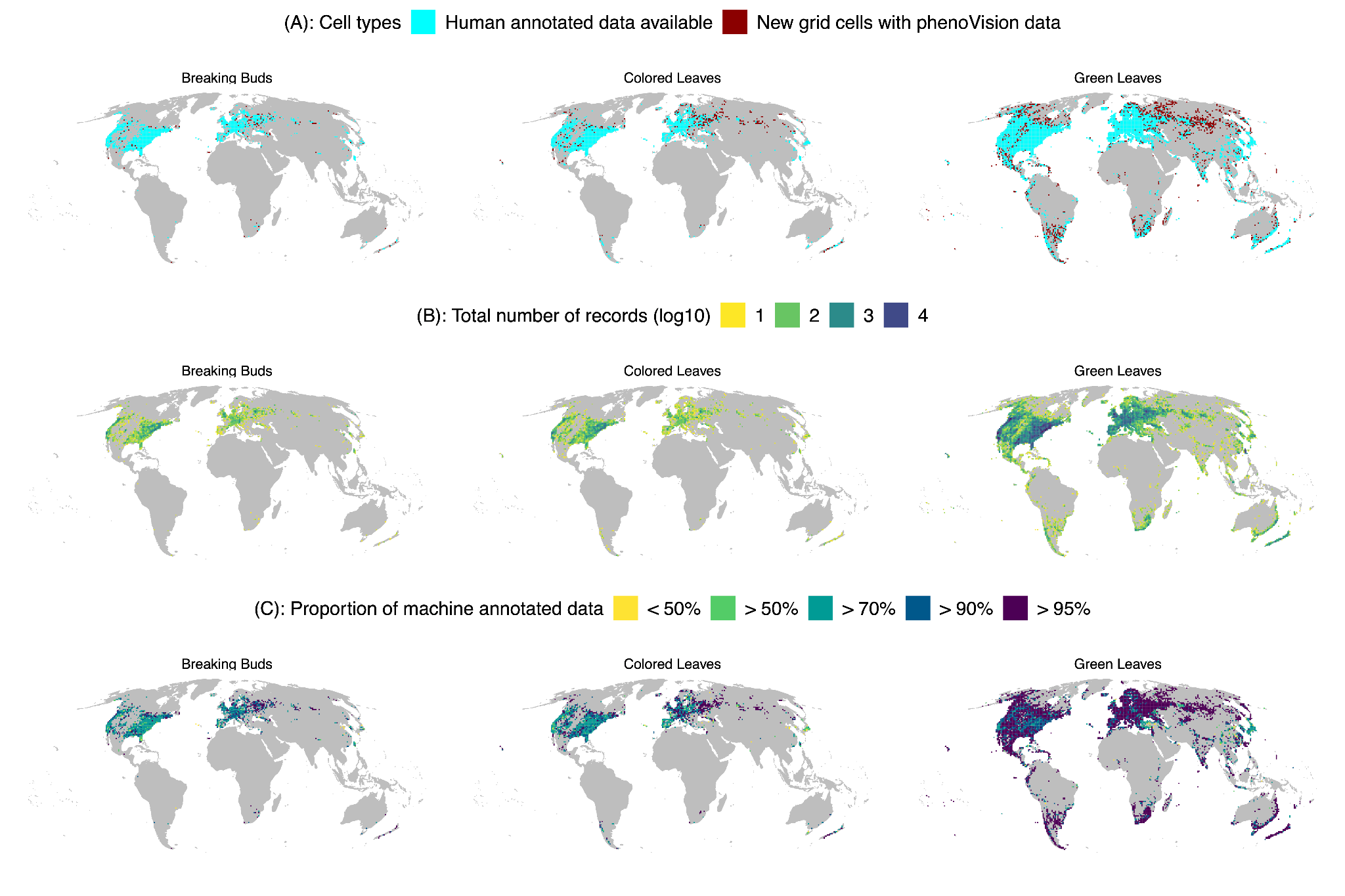

**Figure S4.** Global spatial coverage of breaking leaf bud, colored leaf, and green leaf phenology records at a 100km x 100km equal-area grid cell level. Each colored grid cell represents a cell with at least ten phenophase records; grey cells represent gaps. (A) The spatial distribution of grid cells with at least one manually-annotated phenophase present record (light blue) and new grid cells for which PhenoVision provided machine-annotated data (dark red). (B) The spatial distribution of record counts (log_10_-transformed) for each leaf phenophase generated by PhenoVision. (C) The spatial distribution of the proportion of phenology data generated by PhenoVision.
